# TREM2 Hit Discovery Using TRIC Technology: A Proof-of-Concept High-Throughput Screening Approach

**DOI:** 10.1101/2025.06.03.657564

**Authors:** Natalie Fuchs, Katarzyna Kuncewicz, Farida El Gaamouch, Moustafa T. Gabr

## Abstract

Triggering receptor expressed on myeloid cells 2 (TREM2) is an immunomodulatory receptor implicated in both neurodegenerative diseases and cancer. Depending on the context, TREM2 agonists or inhibitors hold therapeutic potential. To date, the majority of TREM2-targeted strategies have centered on monoclonal antibodies (mAbs), which face limitations such as poor tissue penetration and potential immunogenic side effects. To overcome these challenges and expand the chemical space for TREM2-targeting agents, we developed a high-throughput screening (HTS) platform to identify novel small molecule TREM2 binders. Using temperature-related intensity change (TRIC) technology in a 384-well plate format (NanoTemper Dianthus), we screened two focused compound libraries comprising over 1,200 molecules. From this screen, 18 preliminary hits (1.44% hit rate) were identified and subsequently validated by dose-response binding studies using microscale thermophoresis (MST), yielding four validated hits (0.32% hit rate) with binding affinities in the high to medium micromolar range (e.g., T2337, *K*_D_ = 22.4 μM). The binding of the top hit, T2337, was further validated using surface plasmon resonance (SPR). Additionally, we assessed the functional activity of all four validated hits in a cellular assay measuring TREM2-mediated Syk phosphorylation in HEK293 cells co-expressing human TREM2 and its adaptor protein DAP12. These findings establish a robust and scalable platform for the discovery of small molecule TREM2 modulators and serve as a proof-of-concept for broader HTS campaigns targeting TREM2.

## 1. Introduction

Triggering receptor expressed on myeloid cells 2 (TREM2) is an immunomodulatory receptor expressed on innate immune cells and plays a key role in various diseases, including neurodegenerative disorders and cancers.^1,2^ TREM2 is essential for the activation and survival of myeloid cells, making it a promising therapeutic target for neurodegenerative diseases such as Alzheimer’s disease (AD), as well as for cancer immunotherapy strategies.^1,3^

With regard to neurodegeneration, TREM2 plays a key role in regulating microglial functions such as phagocytosis.^4,5^ Studies suggest that specific TREM2 mutations, such as R47H, may increase the risk of Alzheimer’s disease (AD) by impairing microglial clearance of amyloid plaques and tau tangles.^6-11^ TREM2 promotes different microglial activation states, some of which are beneficial for amyloid plaque clearance and others unfavorable.^10,12^ In addition, TREM2 expression levels and mutations are associated with tau phosphorylation which contributes to the formation of tau tangles.^6-8,13^ The interplay between TREM2 and AD remains multifaceted, as its functional impact appears to depend on both the stage of disease progression and the particular TREM2 variants present.^5,14^ Nevertheless, researchers have explored TREM2 agonists (e.g., monoclonal antibodies(mAbs)) to improve microglial phagocytosis as a potential therapeutic strategy to treat AD.^15^

As a player in the tumor microenvironment (TME), TREM2 is overexpressed in multiple tumors, including lung and ovarian cancer,^3,16^ where it is predominantly expressed on tumor-associated macrophages (TAMs) and myeloid-derived suppressor cells (MDSCs).^17,18^ TREM2 expression is associated with CD8^+^ T cell exhaustion in peripheral tumors, fostering an immunosuppressive TME.^19,20^ Recent studies suggest that TREM2 inhibition or knock-out can improve T cell responses to tumor cells and attenuate tumor growth.^17,20^ Additionally, targeting TREM2 can re-sensitize tumors for anti-PD-1 therapy, emphasizing the potential of a combination therapy.^21,22^ To date, therapeutic approaches targeting TREM2 have focused solely on mAbs which have several drawbacks, including poor tumor penetration and bioavailability, high manufacturing costs and potentially immunogenic side effects.^23,24^

Our goal is to identify small molecule ligands for TREM2 as an alternative to mAbs, aiming to overcome their inherent limitations. Small molecules present a promising alternative to mAbs as their properties can be optimized in terms of bioavailability and tissue penetration.^23^ High throughput screening (HTS) is a powerful tool to find novel small molecule binders with diverse chemical scaffolds.^25^ These screening campaigns can be performed using biophysical platforms, cell-based assays or biochemical assays.^26,27^ Biophysical screening platforms are best suited for the search for new TREM2 binders, as they can easily be scaled up for large chemical libraries and provide reliable results for protein-ligand binding without additional factors (e.g., co-factors for enzymatic reactions, other interfering proteins in cell-based assays).^27,28^ Common HTS screening platforms include surface plasmon resonance (SPR), biolayer interferometry (BLI) or fluorescence-based assays.^27,29,30^

In this study, we established an HTS platform for TREM2 using temperature-related intensity change (TRIC) in a 384-well plate format (NanoTemper Dianthus), a fluorescence-based biophysical technique to determine molecular interactions. We selected two small, focused libraries based on literature ligands (TargetMol Lipid Metabolism library, TargetMol Saccharide and Glycoside Natural Product library), totaling 1,248 compounds. We set up the HTS platform as well as a validation platform with dose-dependent studies using microscale thermophoresis (MST, NanoTemper Monolith) to find new potential small molecule TREM2 binders. We performed orthogonal validation of the top hit compound using surface plasmon resonance (SPR) and tested the ability of the identified hits to modulate TREM2 signaling in vitro.

## 2. Materials and Methods

### 2.1 High-throughput screening with Dianthus

The HTS was performed using a TRIC-based method on Dianthus NT.23Pico (NanoTemper, Munich, Germany). The compound libraries were purchased from TargetMol (Boston, MA, USA), namely “Lipid Metabolism Compound Library” (653 compounds, #L2510) and “Saccharide and Glycoside Natural Product Library” (595 compounds, #L6140). His-tagged TREM2 (SinoBiological, #11084-H08H) was labeled with RED-tris-NTA 2^nd^ generation dye (NanoTemper, #MO-L018) according to the manufacturer’s protocol.

The libraries were transferred from 384-well DMSO stock plates (10 mM) to intermediate plates at a 2-fold ligand concentration (200 μM, PBS pH 7.4, 0.05% Tween20, 2.5% DMSO) using the Integra mini-96 pipetting robot (**Figure 1**).

**Figure 1.**
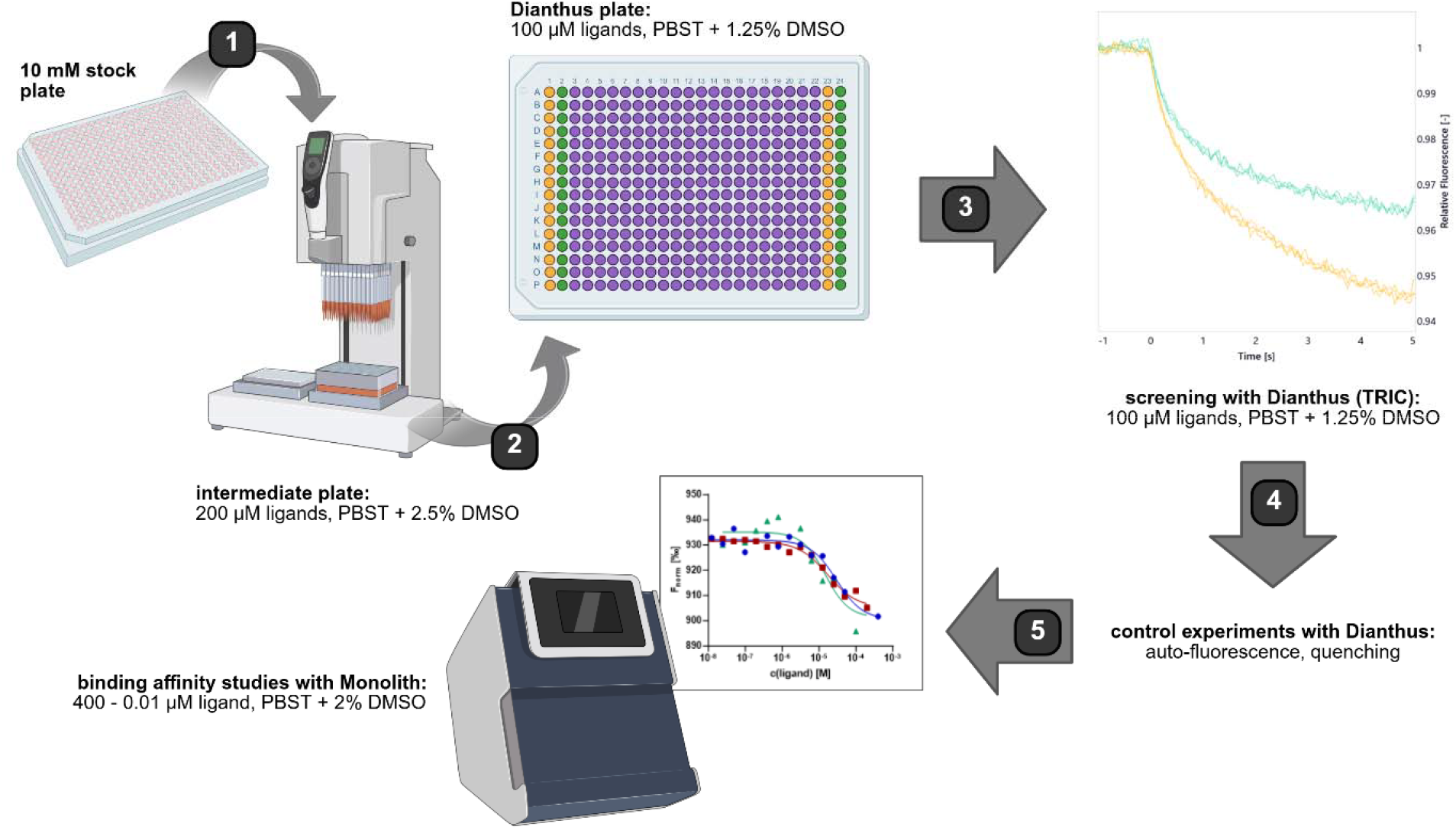
Project workflow. (**1**) Preparation of intermediate plates from library stock using Integra mini-96. (**2**) Dianthus plates were prepared from intermediate plates using Integra mini-96. (**3**) HTS with Dianthus and screening conditions. (**4**) Control experiments with Dianthus to identify false positive hits. (**5**) Hit validation with Monolith NT.115 to determine binding affinity and screening conditions. Figure composed with BioRender.

For the screening in a 384-well plate format, 10 nM labeled His-TREM2 was incubated with 100 μM compound in assay buffer (PBS, pH 7.4, 0.05% Tween20, final DMSO concentration 1.25%) for 10 minutes. Additionally, a negative control (PBS, pH 7.4 0.05% Tween20, 1.25% DMSO) was run in two columns per plate (**Figure 1**). The normalized fluorescence (F_norm_) for all wells was analyzed with GraphPad Prism 10. Compounds with an F_norm_ outside of a 10 standard deviation range from the average control were considered as potential hits. Compounds that were flagged by the instrument’s software (DI.Control) for aggregation or scan anomalies were excluded from the analysis. A more detailed description of the hit selection can be found in the **Supporting Information**.

### 2.2 Control experiments

Control experiments were performed to check the potential hits for assay interference. In a first test, the hit compounds were incubated with assay buffer and the initial fluorescence and TRIC signal were assessed with Dianthus. For a second control experiment, the hit compounds were incubated with RED-tris-NTA 2^nd^ generation dye and their initial fluorescence and TRIC signal were measured using Dianthus (**Figure 1**). Compounds that did not show any significant difference from references (PBS, pH 7.4, 0.05% Tween20, 1.25% DMSO) were considered for further experiments. All remaining hit compounds were verified with a repetition of the single-dose screening at 100 μM (n = 3).

### 2.3 Binding affinity with MST

Binding affinity studies were performed on Monolith NT.115 (NanoTemper, Munich, Germany). Ligands (16 assay points, 400–0.01 μM, 2-fold dilution series) were incubated with 25 nM His-TREM2 labeled with RED-tris-NTA 2^nd^ generation as described above in assay buffer (PBS, pH 7.4, 0.05% Tween20, 2% DMSO) for 10 minutes (**Figure 1**). MST measurements were conducted at 100% laser excitation and medium MST power. *K*_D_ values were determined by the Monolith software (MO.Affinity Analysis) using the F_norm_ values that were converted to fraction bound values by the analysis software. *K*_D_ values present an average from three independent experiments. Graphs for figures were prepared using GraphPad Prism 10.

### 2.4 SPR validation

The binding of T2337 for human TREM2 protein was determined by SPR (Biacore 8K, Cytiva). The biotinylated TREM2 protein was immobilized on SA Sensor Chip (Cytiva). Five concentrations of the compound (120, 60, 30, 15 and 7.5 μM) were prepared in PBS-P buffer + 3% DMSO (Cytiva) and injected over the prepared surface of the SA sensor chip. The runs were conducted at 25°C employing the following parameters: flow rate of 30 μl/min, contact time of 120 s, and dissociation time of 300 s. After each analysis an additional wash with 50% DMSO solution was performed. The results are presented as sensorgrams obtained after subtraction of the background response signal from a reference flow cell and from a control experiment with buffer injection. The obtained data were analyzed using BiacoreTM Insight Evaluation Software (Cytiva). Single cycle kinetic analysis was performed using a 1:1 reaction model, giving the best fitting of experimental and theoretical data.

### 2.5 In vitro TREM2 activation

Phospho-AlphaLISA assay measures a protein target when phosphorylated at a specific residue. The assay uses two antibodies which recognize the phospho epitope and a distal epitope on the targeted protein. In the presence of phosphorylated protein, luminescent Alpha signal is generated. The amount of light emission is directly proportional to the quantity of phosphoprotein present in the sample. Briefly, HEK–hTREM2/DAP12 cells were seeded at 50,000 cells per well in 96-well plate, in a final volume of 100 μl of growth media composed of DMEM (Gibco) supplemented with 10% FBS (Gibco) and incubated for 24 h at 37 °C, 5% CO_2_. The four tested compounds were diluted in media at a concentration of 100 μM, growth media was manually removed from the wells and subsequently replaced with the test compounds containing media. Control wells were treated with the vehicle buffer used to solubilize the several compounds. After 1h, the media was gently removed, lysis buffer was added and after complete lysis, AlphaLISA assay was performed according to manufacturer instructions (Revvity, USA).

## 3. Results and Discussion

In this focused HTS approach, we screened at total of 1,248 compounds from two libraries (TargetMol Lipid Metabolism Compound library and TargetMol Saccharide and Glycoside Natural Product library, **Figure 2**). We selected both libraries based on literature-known, mainly endogenous TREM2 ligands, which are often described as anionic and/or lipidic compounds (e.g., phospholipids or glycosides and saccharides).^1,31-34^

**Figure 2.**
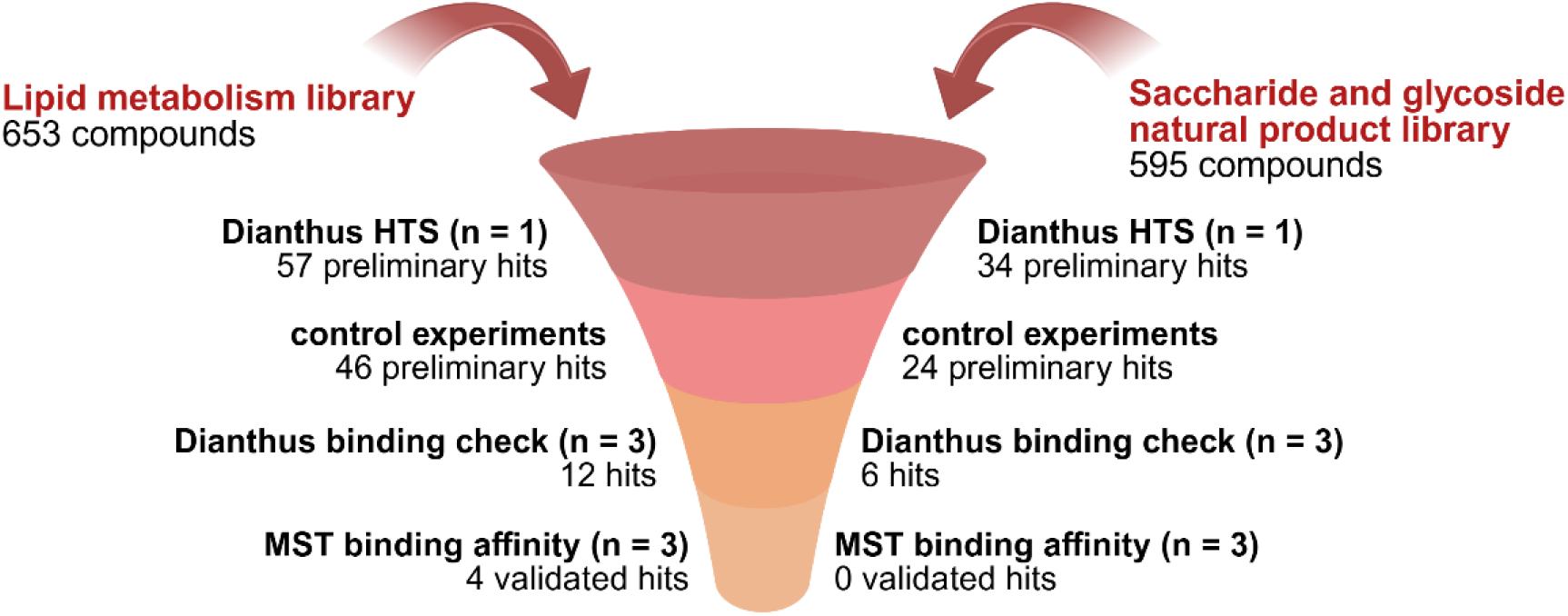
Screened libraries and steps during the HTS and validation process. Figure composed with BioRender.

### 3.1 Single-dose screening with Dianthus

The initial single-dose screening (n = 1) resulted in 57 hits for the Lipid Metabolism Compound library (preliminary hit rate: 8.7%, **Figure 3A**) and 34 hits for the Saccharide and Glycoside Natural Product library (preliminary hit rate: 5.7%, **Figure 3B**).

**Figure 3.**
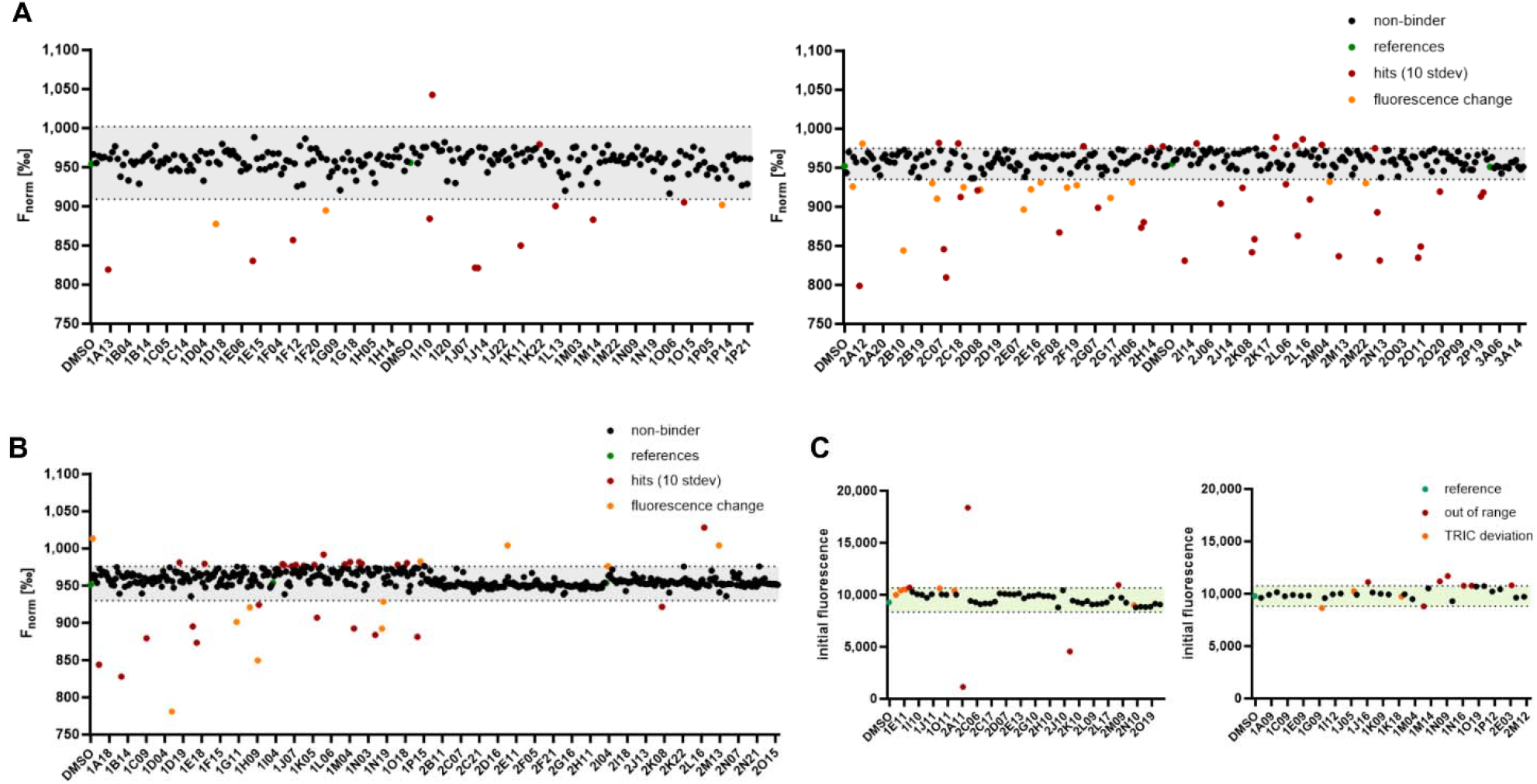
Dianthus screening results. (**A**) TargetMol Lipid Metabolism library, plate 1 (left) and 2 (right). (B) TargetMol Saccharide and Glycoside Natural Product library, both plates combined in one plot. (C) Results of the control experiments for hits from the Lipid Metabolism (left) and Saccharide and Glycoside Natural Product libraries (right). All graphs were created with GraphPad Prism 10 and the figure was composed with BioRender.

Since some compounds were out of range for the initial fluorescence (more than 20% difference to DMSO reference), we conducted control experiments for all hits to exclude false positives. The controls were run with assay buffer and RED-Tris NTA dye to see if the compounds interfere with the dye rather than the protein. This reduced the hit number to 46 for the Lipid Metabolism library (preliminary hit rate: 7.0%, **Figure 3C**) and 24 hits for the Saccharide and Glycoside library (preliminary hit rate: 4.0%, **Figure 3C**).

To further narrow down the number of hits, we repeated the binding check for all hits that did not interfere with the assay conditions (100 μM, n = 3). After control experiments and the additional single-dose binding checks, the final number of hits was 12 for the Lipid Metabolism library (hit rate: 1.84%) and 6 for the Saccharide and Glycoside Natural Product library (hit rate: 1.01%), resulting in an overall hit rate of 1.44%.

Based on our findings, TRIC is a suitable technique for HTS of small molecule libraries targeting TREM2. However, the platform does present certain limitations that should be considered. The required fluorescent protein labeling can lead to assay interferences with small molecules that have fluorescent or quenching properties, yielding false positive hits (e.g., 23% out of the initial hits in this study). Therefore, careful selection of compound libraries is essential to minimize the risk of assay interference and ensure reliable screening results. The Dianthus software flags compounds whose fluorescence is not within 20% of the controls. Nevertheless, testing all hits in control experiments is critical to exclude false positives, even if they are not initially indicated by the software.

Overall, the TRIC assay offers straightforward setup and optimization, making it an appealing alternative to SPR for screening applications. Another advantage is the immobilization-free approach, since immobilizing the protein could potentially interfere with its binding site. In our setup, we use site-directed labeling by attaching the dye to the His tag as an attempt to minimize binding site interference. Additionally, our TRIC screening platform enables rapid high-throughput screenings with low sample consumption, since only small amounts of protein are required to generate sufficient signal, and a 384-well plate can be screened within 30 minutes. Short assay times are beneficial as they reduce the risk of protein degradation and maintain sample integrity. These results are encouraging for future high-throughput screenings for TREM2 on an even larger scale.

### 3.2 Hit validation with Monolith

To validate the Dianthus hits, we tested the 18 remaining compounds for binding affinity with MST. Longer laser on times in the Monolith make it a more reliable instrument for dose-dependent studies than Dianthus. However, the technique is based on the same principle as our high-throughput screening platform. Therefore, it is crucial to perform all control experiments to avoid false positive hits. We were able to exclude 14 out of 18 compounds that did not show dose-dependent binding, possibly due to non-specific binding at higher doses. Quality controls based on the MST traces also helped to exclude potential aggregators.

Ultimately, only four compounds from the Lipid Metabolism Compound library emerged to bind TREM2 in a dose-dependent manner, giving a final rate of 0.32% for validated hits. The *K*_D_ values are in the high and medium micromolar range, with compound T2337 (BMS-303141, assay well 1G04) as the top hit with a *K*_D_ of 22.4 ± 10.2 μM (**Figure 4**). The top compound was developed by Li and co-workers as a potential ATP-citrate lyase inhibitor for hypolipidemic intervention.^35^ More information on all hits can be found in the **Supporting Information**. Overall, the hit validation with MST showed the potential to find dose-dependent binders with our HTS assay setup.

**Figure 4.**
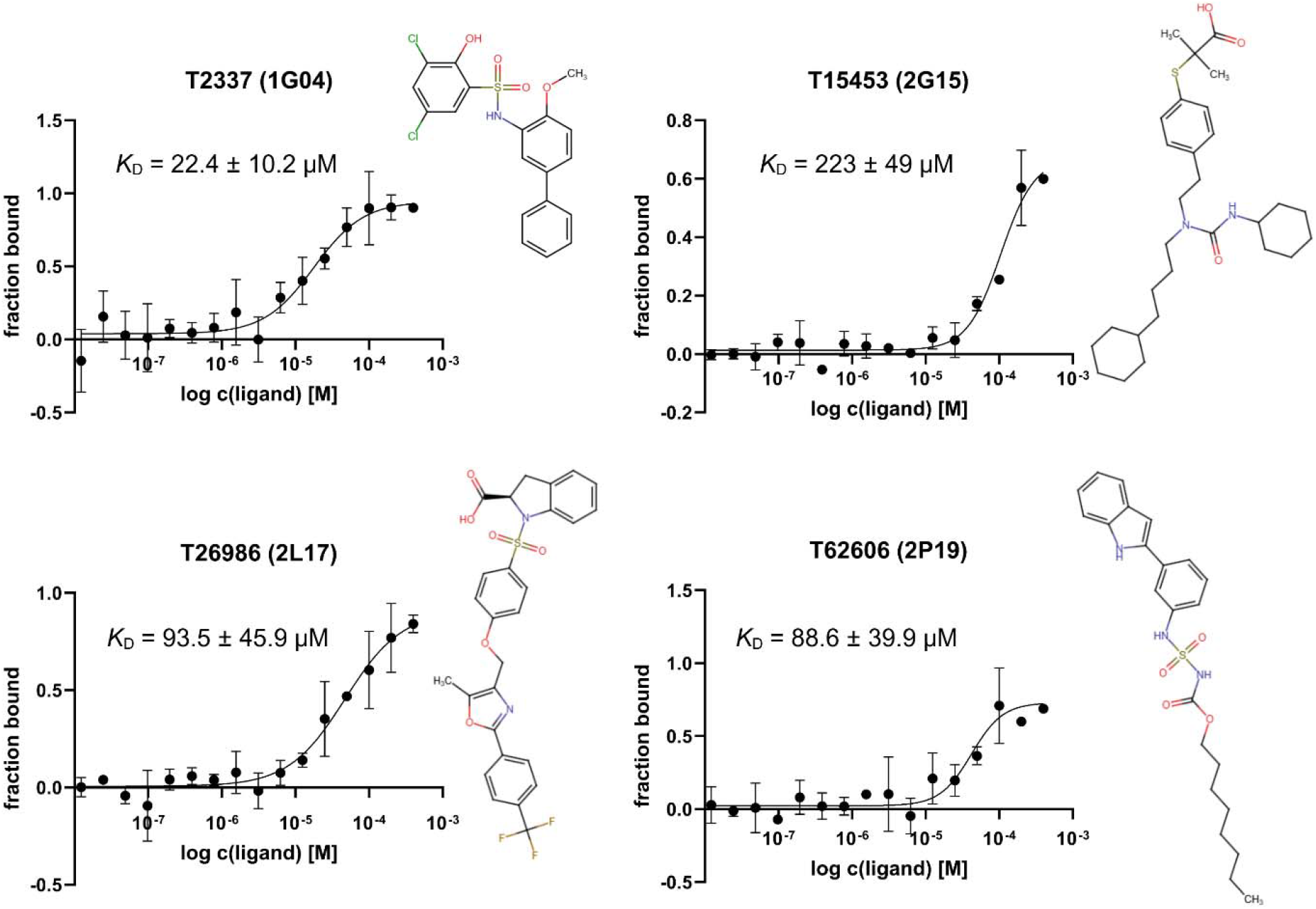
Binding affinity results for four validated hits shown as fraction bound vs log c plots. The binding affinities are in the high to medium micromolar range. All graphs were created with GraphPad Prism 10 and the figure was composed in BioRender.

### 3.3 SPR validation

To independently confirm the interaction between T2337 and TREM2, we employed SPR as an orthogonal method. SPR is a sensitive, real-time, label-free technique widely used to study biomolecular interactions, making it a suitable complement to MST. In this validation experiment, TREM2 was immobilized on the sensor chip, and varying concentrations of T2337 were injected across the surface. The resulting sensorgram (Figure 5) showed a clear, concentration-dependent increase in binding signal, consistent with a specific interaction between T2337 and TREM2. This SPR-based validation supports the MST findings and reinforces the conclusion that T2337 directly binds to TREM2.

**Figure 5.**
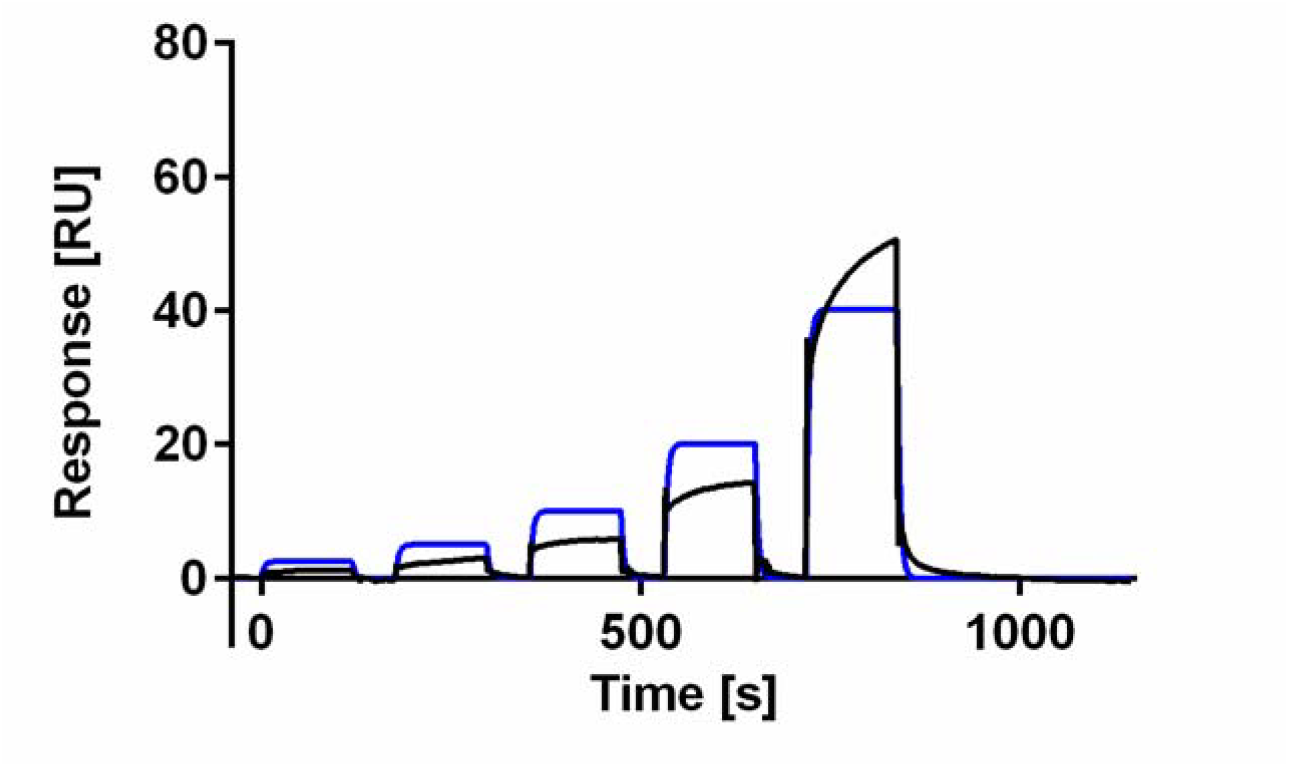
Single-cycle kinetic sensorgrams of T2337 (120, 60, 30, 15, and 7.5 μM) interacting with the TREM2 protein using SPR. Black lines represent the experimental data, while blue lines show the 1:1 kinetic binding model fit.

### 3.4. In vitro evaluation

The four hit compounds, identified from the screening assays, were tested for their ability to activate TREM2 signaling in vitro using HEK293 cells engineered to express human TREM2 and its adaptor protein DAP12. Activation of the TREM2/DAP12 complex triggers phosphorylation of spleen tyrosine kinase (SYK), leading to downstream signaling pathways involved in microglial function. SYK activation was evaluated using a commercial AlphaLISA kit to detect phosphorylation at tyrosine residues 525/526. Fluorescence was measured with a TECAN SPARK plate reader under AlphaLISA settings. As illustrated in Figure 6, T2337 (1G04), the top-performing compound from this study, enhanced SYK phosphorylation relative to the control, whereas the other compound showed minimal impact on TREM2 activation. Although not highly potent, T2337 (1G04) emerges as a promising starting point for hit-to-lead optimization to developed potential TREM2-targeted therapies for AD.

**Figure 6.**
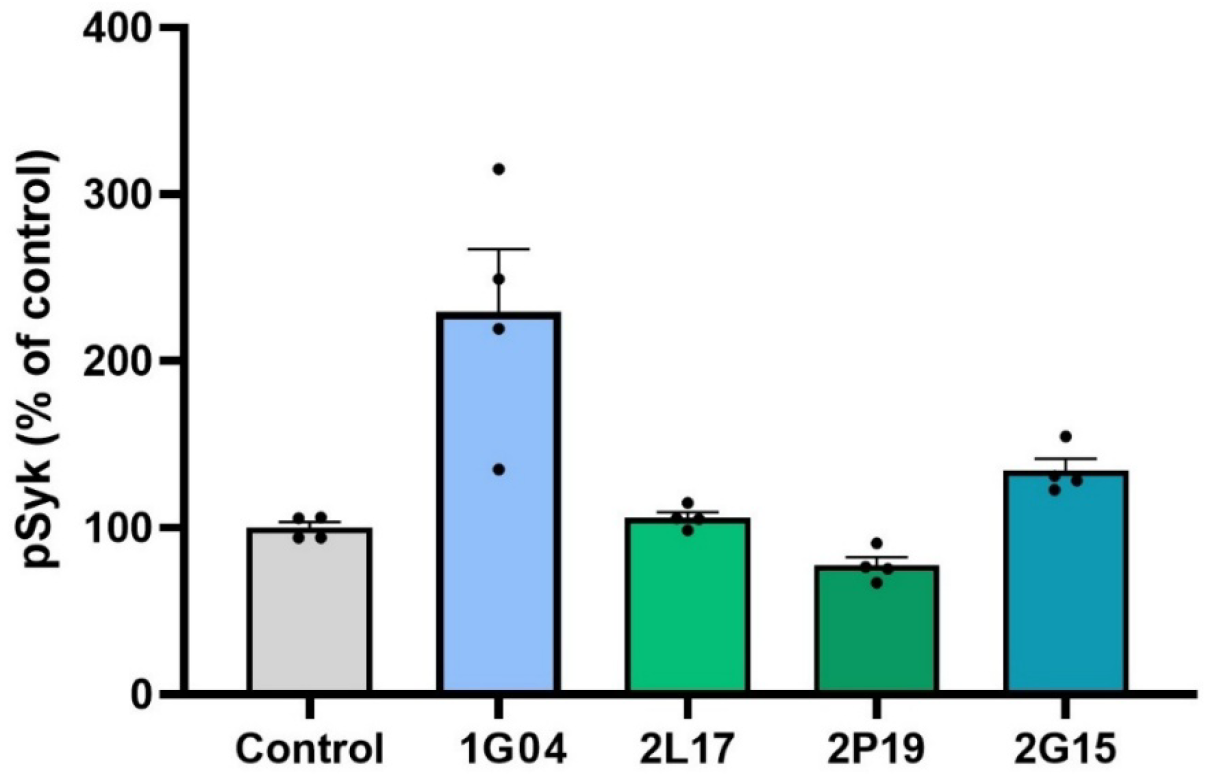
Histogram representing the quantification of phospho-Syk levels, measured using AlphaLisa technique, in untreated condition or after treatment with T2337 (1G04), TL26989 (2L17), T62606 (2P19), and T15453 (2G15) (100 μM-1 h). Phospho-Syk changes in treated conditions are expressed as percentage of control (N=4).

In summary, we employed TRIC to establish a platform that enables screening chemical libraries of small molecules as potential TREM2 ligands on a large scale. We validated our platform using two focused libraries totaling more than 1,200 compounds, which resulted in four true hits (final hit rate 0.32%) that display dose-dependent binding with MST and SPR. TRIC proved well-suited for rapid HTS due to its low sample requirements—only minimal protein amounts are needed to achieve robust signals, and a full plate can be screened in under 30 minutes. The short assay duration also helps preserve protein integrity by minimizing degradation over time. Implementing our TRIC assay in larger screening campaigns in the future will enable the discovery of novel small molecule TREM2 hits with diverse chemical scaffolds. This expanded effort has the potential to uncover new chemotypes for further optimization and functional validation, ultimately contributing to the development of TREM2-targeted therapeutic agents.

## CRediT Authorship Contribution Statement

Natalie Fuchs: Conceptualization, Methodology, Validation, Formal Analysis, Investigation, Data Curation, Writing – Original Draft, Visualization. Katarzyna Kuncewicz: Methodology, Formal Analysis, Data Curation. Farida El Gaamouch: Methodology, Formal Analysis, Data Curation. Moustafa T. Gabr: Writing - Review & Editing, Funding Acquisition.

## Acknowledgments

This work was supported by the National Institutes on Aging under grant number R01AG083512 (PI: Gabr). We would like to thank the Fisher Drug Discovery Resource Center of Rockefeller University (RRID:SCR_020985) for providing access to the Nanotemper Dianthus NT.23 Pico, Nanotemper Monolith NT.115, and Cytiva Biacore 8K instruments.

## Data Availability Statement

The data that support the findings of this study are available in the Supporting Information.

