## Supporting Information for "TREM2 Hit Discovery Using TRIC Technology: A Proof-of-Concept High-Throughput Screening Approach"

### Hit selection

#### Initial single-dose screening

The average F_norm_ was calculated for all references (32 replicates per plate, 16 replicates per laser) followed by a 10 standard deviation range. Since we observed small deviations between the top and bottom laser (top laser: rows A–H, bottom laser: rows I–P), we decided to compare the F_norm_ of the tested compounds to that of the references that were measured with the same laser (plate layout see **Figure 1**). Compounds outside this range were considered potential hits (**Figure 2** and **Figure 3**).

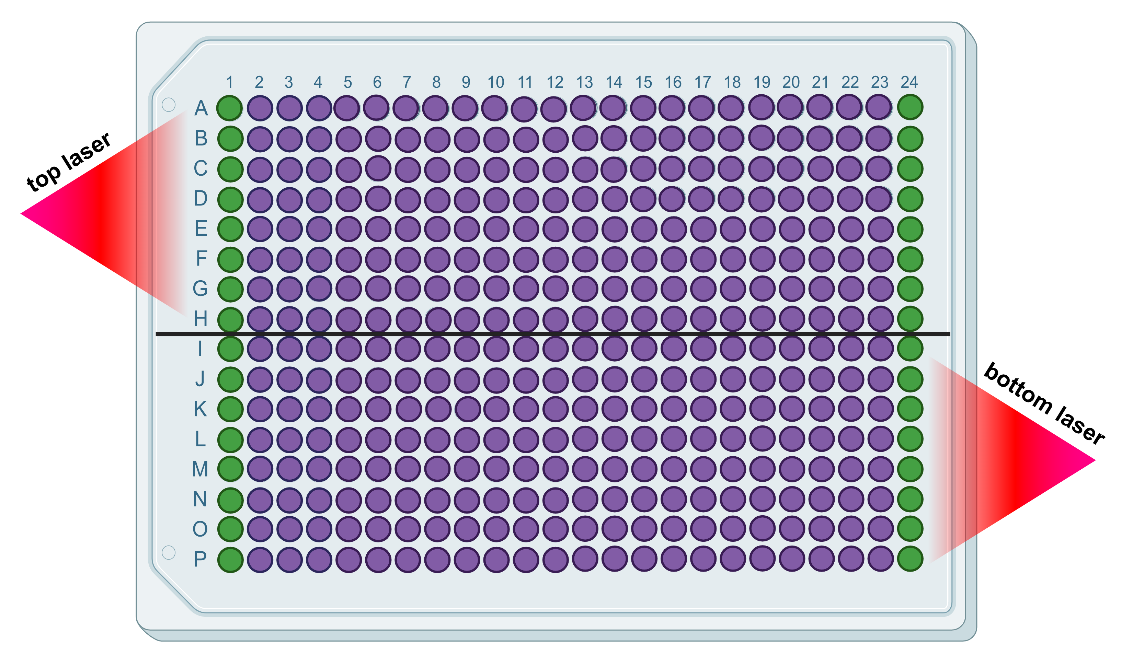

**Figure 1.** Plate layout including location of two lasers. Green wells represent the references (DMSO in PBST), purple wells the samples. The top laser read wells A1–H24, the bottom laser wells I1–P24. Figure created with BioRender.

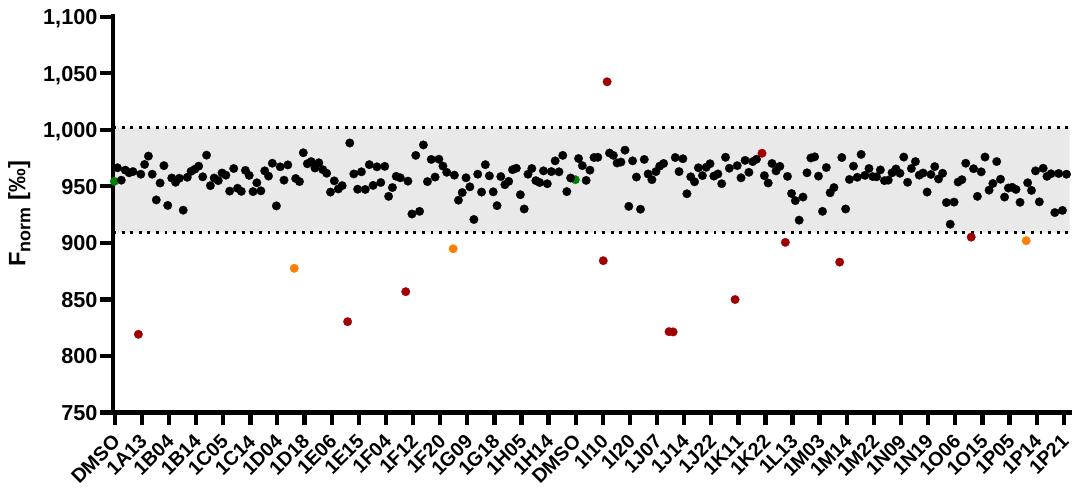

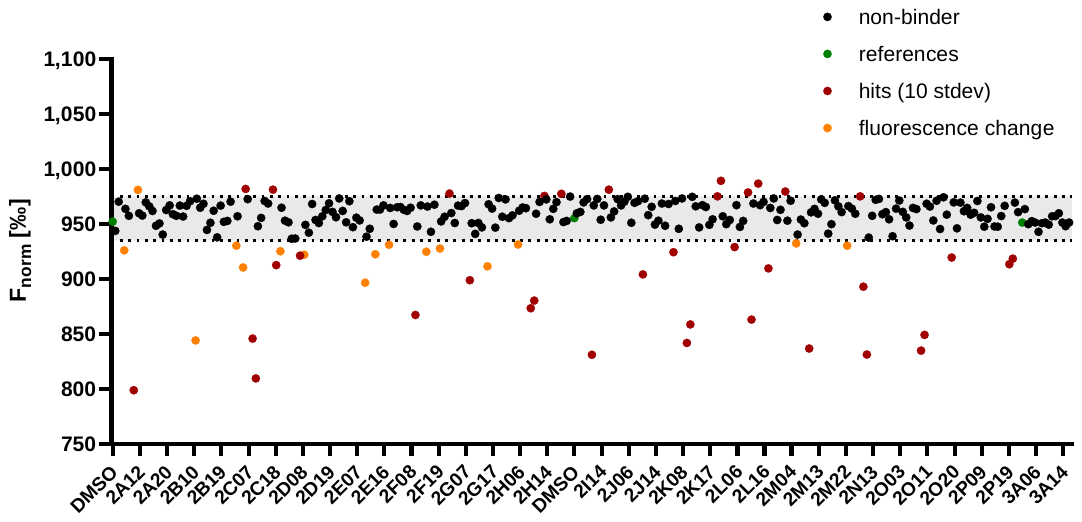

**Figure 2.** Screening results for plates 1–3 of the TargetMol Lipid Metabolism Compound library at 100 µM ligand concentration (n = 1). Reference (DMSO in PBST) shown in green as mean of 16 replicates per laser. Orange dots indicate a change of initial fluorescence compared to references. Hits without fluorescence change are shown in red.

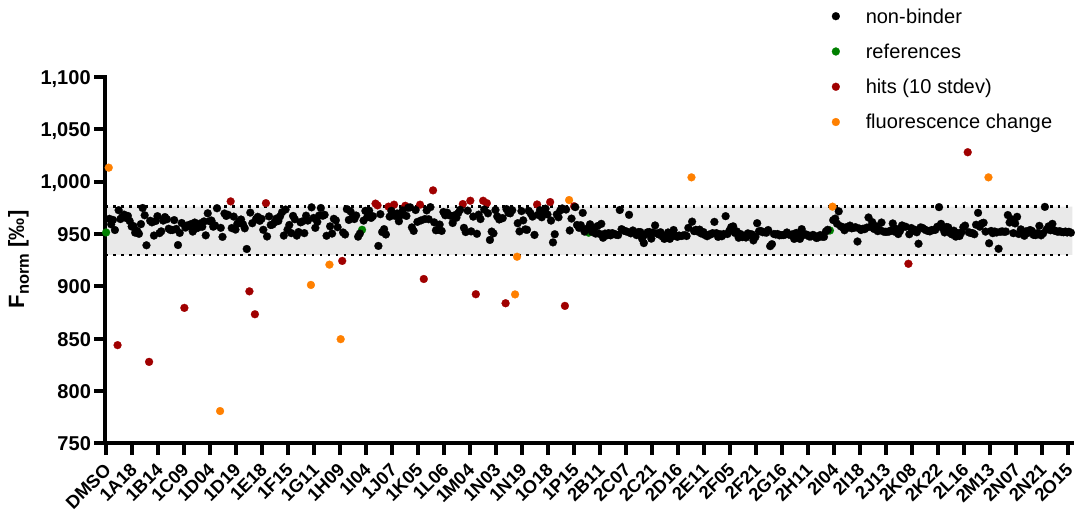

**Figure 3.** Screening results for plates 1–2 of the TargetMol Saccharide and Glycoside Natural Product library at 100 µM ligand concentration (n = 1). Reference (DMSO in PBST) shown in green as mean of 16 replicates per laser. Orange dots indicate a change of initial fluorescence compared to references. Hits without fluorescence change are shown in red.

#### Control experiments

**Table 1.** Results of control experiments for TargetMol Lipid Metabolism library hits. DMSO in PBST as reference, compounds that deviated in orange, compounds without deviations in green.

| **sample** | **initial fluorescence** | **out of range** | **TRIC trace deviation** |
| --- | --- | --- | --- |
| **DMSO** | **9273** |  |  |
| 1A11 |  |  | 9996 |
| 1D09 |  |  | 10407 |
| 1E11 |  |  | 10529 |
| 1F10 |  | 10703 |  |
| 1G04 | 10289 |  |  |
| 1I10 | 10066 |  |  |
| 1I13 | 10014 |  |  |
| 1J10 | 9725 |  |  |
| 1J11 | 10062 |  |  |
| 1K10 |  |  | 10620 |
| 1M10 | 10044 |  |  |
| 1O11 | 10011 |  |  |
| 1P10 |  |  | 10401 |
| 2A10 | 10016 |  |  |
| 2A11 |  | 1156 |  |
| 2B10 |  | 18386 |  |
| 2C05 | 9414 |  |  |
| 2C06 | 9295 |  |  |
| 2C09 | 9092 |  |  |
| 2C10 | 9181 |  |  |
| 2C17 | 9167 |  |  |
| 2C18 | 9355 |  |  |
| 2C19 | 10136 |  |  |
| 2D07 | 10087 |  |  |
| 2D08 | 10060 |  |  |
| 2E09 | 10035 |  |  |
| 2E13 | 10134 |  |  |
| 2F10 | 9674 |  |  |
| 2F14 | 9887 |  |  |
| 2G10 | 9899 |  |  |
| 2G15 | 10032 |  |  |
| 2H09 | 9879 |  |  |
| 2H10 | 9892 |  |  |
| 2I10 | 9797 |  |  |
| 2I19 | 8787 |  |  |
| 2J10 | 10431 |  |  |
| 2J20 |  | 4553 |  |
| 2K09 | 9447 |  |  |
| 2K10 | 9313 |  |  |
| 2K21 | 9177 |  |  |
| 2L05 | 9401 |  |  |
| 2L09 | 9072 |  |  |
| 2L10 | 9115 |  |  |
| 2L13 | 9150 |  |  |
| 2L17 | 9278 |  |  |
| 2L22 | 9753 |  |  |
| 2M05 |  | 10941 |  |
| 2M09 | 9722 |  |  |
| 2M22 | 9256 |  |  |
| 2N09 |  |  | 8994 |
| 2N10 | 8788 |  |  |
| 2O09 | 8852 |  |  |
| 2O10 | 8837 |  |  |
| 2O19 | 8857 |  |  |
| 2P19 | 9152 |  |  |
| 2P20 | 9064 |  |  |

**Table 2.** Results of control experiments for TargetMol Saccharide and Glycoside library hits. DMSO in PBST as reference, compounds that deviated in orange, compounds without deviations in green.

| **sample** | **initial fluorescence** | **out of range** | **TRIC trace deviation** |
| --- | --- | --- | --- |
| **DMSO** | **9778** |  |  |
| 1A04 | 9618 |  |  |
| 1A09 | 9908 |  |  |
| 1B09 | 10144 |  |  |
| 1C09 | 9774 |  |  |
| 1D09 | 9888 |  |  |
| 1E09 | 9812 |  |  |
| 1E12 | 9831 |  |  |
| 1G09 | 8640 |  |  |
| 1H09 | 9585 |  |  |
| 1I12 | 9949 |  |  |
| 1I14 | 10028 |  |  |
| 1J05 |  |  | 10248 |
| 1J08 | 9912 |  |  |
| 1J16 |  | 11108 |  |
| 1K06 | 10117 |  |  |
| 1K09 | 9987 |  |  |
| 1K17 | 9938 |  |  |
| 1K18 |  |  | 9737 |
| 1L20 | 9946 |  |  |
| 1M04 | 9486 |  |  |
| 1M09 |  | 8800 |  |
| 1M14 | 10510 |  |  |
| 1M18 |  | 11179 |  |
| 1N09 |  | 11685 |  |
| 1N15 | 9288 |  |  |
| 1N16 |  | 10769 |  |
| 1O11 |  | 10780 |  |
| 1O19 | 10691 |  |  |
| 1P09 | 10742 |  |  |
| 1P12 | 10215 |  |  |
| 1P15 | 10427 |  |  |
| 2E03 |  | 10801 |  |
| 2L18 | 9625 |  |  |
| 2M12 | 9717 |  |  |

### List of hit compounds

#### TargetMol Lipid Metabolism Compound library

**Table 3.** Hits for Lipid Metabolism library after single-dose screening (n = 3) and control experiments.

| plate | well | cpd ID | synonym | MW [g/mol] | structure | *K*_D_ [µM] |
| --- | --- | --- | --- | --- | --- | --- |
| 1 | G04 | T2337 | BMS-303141^1^ | 424.3 | 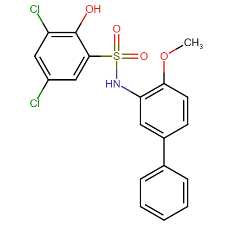 | 22.4 |
| 2 | C18 | TQ0105 | CAY10650^2^ | 489.0 | 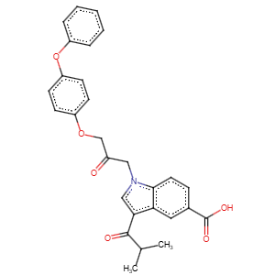 | - |
| 2 | C19 | T8320 | J14^3^ | 517.0 | 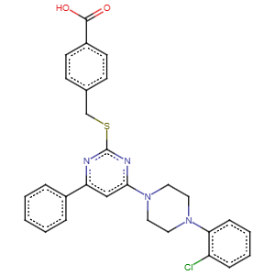 | - |
| 2 | D07 | T36841 | IPI-9119^4^ | 495.4 | 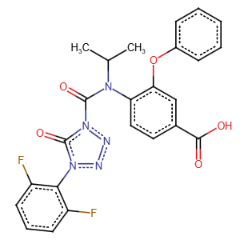 | - |
| 2 | D08 | T27303 | farglitazar^5^ | 546.6 | 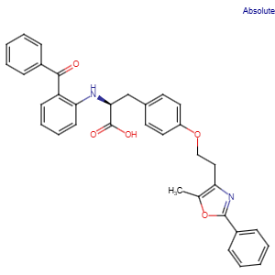 | - |
| 2 | E13 | T8471 | vonafexor^6^ | 489.8 | 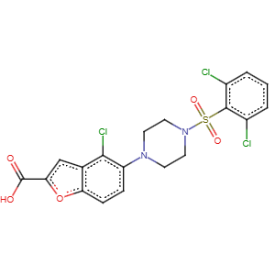 | - |
| 2 | F14 | T16861 | SB 204990^7^ | 389.3 | 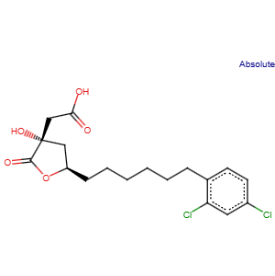 | - |
| 2 | G15 | T15453 | GW7647^8^ | 502.8 | 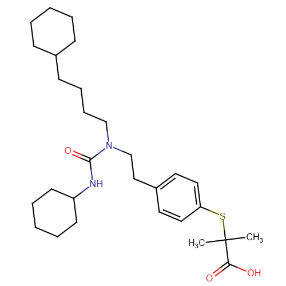 | 223 |
| 2 | L05 | T12834 | saroglitazar magnesium^9^ | 901.4 | 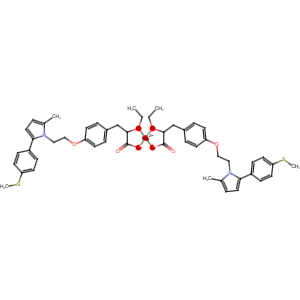 | - |
| 2 | L17 | T26986 | cevoglitazar^10^ | 558.5 | 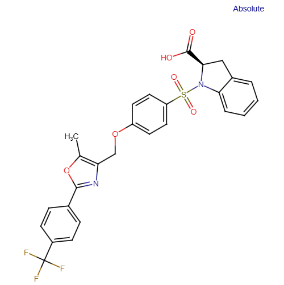 | 93.5 |
| 2 | M22 | T13803 | *N*-oleoyl glycine^11^ | 339.5 | 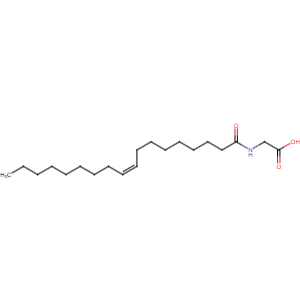 | - |
| 2 | P19 | T62606 | ALOX15-IN-2^12^ | 443.6 | 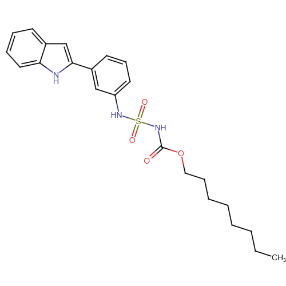 | 88.6 |

#### TargetMol Saccharide and Glycoside Natural Product library

**Table 4.** Hits for Saccharide and Glycoside library after single-dose screening (n = 3) and control experiments.

| plate | well | cpd ID | synonym | MW [g/mol] | structure | *K*_D_ [µM] |
| --- | --- | --- | --- | --- | --- | --- |
| 1 | D09 | T3794 | penta-*O*-galloyl-β-d-glucose^13^ | 940.7 | 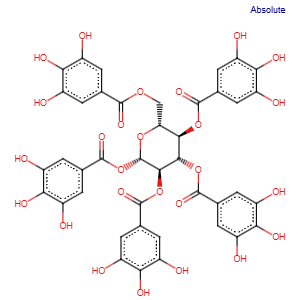 | - |
| 1 | G09 | T2964 | *Astragalus* polyphenols | 406.4 | 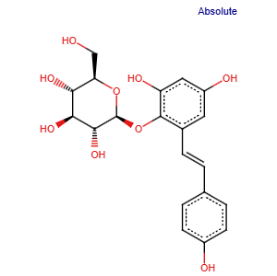 | - |
| 1 | H09 | T2S0389 | emodin-1-*O*-β-d-gluco-pyranoside^14^ | 432.4 | 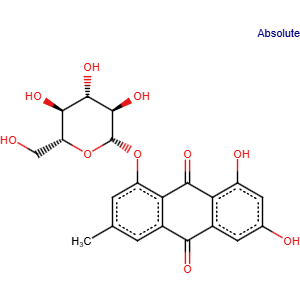 | - |
| 1 | N15 | T6S2099 | geraniin^15^ | 952.6 | 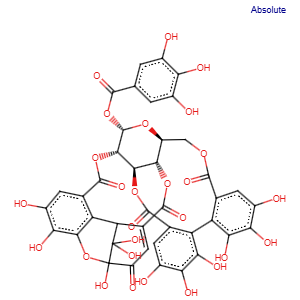 | - |
| 1 | O19 | T2891 | ammonium glycyrrhizinate | 840.0 | 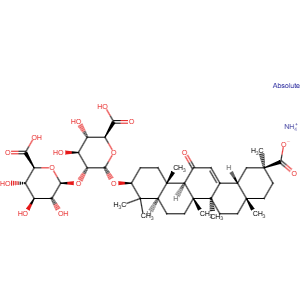 | - |
| 1 | P09 | T2S0690 | ecliptasaponin A^16^ | 634.8 | 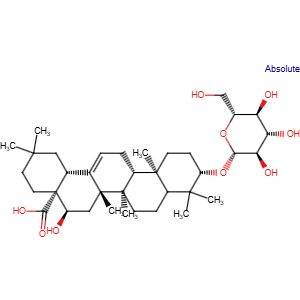 | - |
